# Integrating the MARTINI2 Coarse-Grained Force Field into HADDOCK3 for Faster Modelling of Large Biomolecular Complexes

**DOI:** 10.64898/2026.04.25.720800

**Authors:** Raphaelle Versini, Victor Reys, Anna Kravchenko, Rodrigo V. Honorato, Alexandre M. J. J. Bonvin

## Abstract

The integration of coarse-grained (CG) approaches into docking workflows offers a powerful strategy for modelling large biomolecular assemblies with reduced computational costs. We present here the implementation of the MARTINI2 coarse-grained force field into the HADDOCK3 integrative modelling platform. This development enables the use of the CG representations and parameters within HADDOCK3 for efficient sampling and scoring of large protein–protein complexes. The implementation takes advantage of the modular and flexible architecture of HADDOCK3, allowing a seamless combination of MARTINI2 representation with the various modules. Conversion from and to all-atom models is integrated into the coarse-grained modelling workflow. The performance of the protocol is first assessed on protein-protein and protein-DNA benchmarks and then illustrated on a few representative large-scale systems, demonstrating a significant reduction in computational costs while maintaining biologically relevant accuracy.

## INTRODUCTION

Biomolecular interactions govern all essential processes within a living organism, with proteins being key components of those molecular processes. Structural characterization of these interactions remains a challenge, as their often transient nature and other experimental factors can limit the acquisition of reliable structural information^1,2^.

Computational modelling of protein structures and protein-protein interactions has made a lot of progress in the last 10 years. In fact, artificial intelligence (AI) is now able to predict protein-protein interactions quite accurately^3–6^, as well as interactions involving nucleic acids, lipids, small molecules and post-translational modifications^7–9^. However, classical physics-based docking approaches, such as HADDOCK^10,11^, remain relevant and complementary to machine learning architectures, as exemplified by antibody-antigen complexes, which can be reliably modeled by a combination of AI and physics-based docking methods using interaction restraints to drive the docking ^12–14^, or protein-glycan complexes^15^ for which current AI models currently lack support.

Although docking is efficient for small- and medium-sized proteins, its application to large biological systems faces substantial computational demands to sample complex conformational landscapes, while AI-based approaches are limited by the maximum number of residues they can effectively handle. Coarse-grained (CG) models can mitigate the limitations of physics-based methods by associating several heavy atoms into one larger particle or bead^16–18^, which effectively reduces the number of particles to take into account in the simulations. Several docking programs use CG representations to reduce computational costs, enabling efficient exploration of large conformational spaces, and smoothing the interaction energy landscape. For instance, ATTRACT uses reduced protein models to perform systematic docking minimization across thousand initial structures within a reasonable computing time^19^. The MARTINI coarse-grained model is currently among the most widely used frameworks for CG simulations; it was first developed for lipids^20^ and later extended to include proteins^21^ and DNA^22^. The MARTINI mapping philosophy is to create one CG particle or bead out of four heavy atoms. This mapping is adapted when atoms are involved in long side chains and cycles, as well as DNA bases. It can represent a wide range of molecular types (high transferability) and enables straightforward back-mapping to atomistic resolution, making it particularly well-suited for use in HADDOCK for integrative modelling applications.

The MARTINI2.2 version was previously implemented in HADDOCK2.4 for both proteins^23^ and DNA^24^. The main docking steps—rigid-body docking and semi-flexible refinement—were performed at the CG level, followed by back-mapping to all atoms (AA) resolution prior to the final energy minimization. The implementation maintained the overall docking performance, while reducing the computational costs, allowing for the modelling of larger systems and/or faster runs.

When redesigning HADDOCK into a modular version^11^, moving away from the rigid workflow with pre-defined and parametrizable steps into a modular platform enabling the definition of custom workflows, its coarse-graining feature was deprioritized considering the large scope of the redesign. Here, we describe the implementation of coarse-graining with the MARTINI2.2^21^ force-field into the latest version of our docking software, HADDOCK3^11^. We added two new modules *topocg* and *cgtoaa* that take care of the atomistic to coarse-grained mapping and vice-versa, a task that was only automated in the web implementation of HADDOCK2.4^10^, while the command-line version required providing both atomistic and coarse-grained models as input. We show on protein-protein and protein-nucleic acid benchmark datasets that HADDOCK3 CG can maintain the success rate obtained with all-atom models while reducing the simulation time. We further illustrate HADDOCK3’s CG usage with three different cases: (1) the human liver phosphofructokinase-1 filament in the R-state conformation^25^, (2) the heptameric KaiC:KaiB 1:6 complex^26,27^, and (3) the PRC1-nucleosome core particle complex^28^.

## MATERIALS AND METHODS

### Integration of the MARTINI2 Coarse-Grained Force Field into HADDOCK3

The implementation of MARTINI2.2^21,22^ into HADDOCK3^11^ builds upon the previous implementation in HADDOCK2.4^23,24^. The converted MARTINI topology and parameter files compatible with the HADDOCK engine CNS (Crystallography and NMR System)^29^, built for HADDOCK2.4^10^ were used for this work, including the MARTINI2^22^ extension to DNA.

Two new modules were created to enable coarse-grained docking. The *topocg* module handles the mapping of the all-atom input structures into their coarse-grained representation and prepares the corresponding CG topology files. This module can only be used after the *topoaa* module, which creates the AA topologies and builds any missing atoms. The refinement module *cgtoaa* performs the reverse operation, i.e., the back-mapping of the CG structures into their AA representation using distance restraints (see below).

The MARTINI^21,22^ mapping philosophy was kept in the *topocg* module, following a four-to-one (4:1) rule for the amino acids’ backbone, meaning all four heavy atoms (N, C_α_, C, O) are represented by a single particle (bead) placed at their center of mass. It was derived from the original Martinize script^21^. The conversion rule is adapted for side chains, and ranges from 4:1 to 2:1 mapping, as well as “small” beads in rings (HIS, PHE, TYR, TRP). As the backbone bead parameters depend on the secondary structure of the molecule, we are using DSSP to determine and encode the secondary structure into the B-factor field of the AA models. The proper parameters are then automatically selected for each backbone bead. If no secondary structure is found, random coil parameters are applied instead (which is also the behavior in case DSSP is not installed). Note that the definition of the secondary structure in CG representation is not a requirement to maintain it in HADDOCK, as most of the structure is kept rigid during the semi-flexible refinement stage. The backbone of DNA is represented by 3 beads, one for the phosphate group (4:1) and two for the sugar (3:1). Bases are mapped into three and four beads for pyrimidines and purines, respectively. The MARTINI2 force-field considers four types of interactions for proteins^21^: polar (P), nonpolar (N), apolar (C), and charged (Q). For nucleic acids, eight beads are added to mimic hydrogen bonding formed between complementary nucleotide base pairs^22^, contributing to the stabilization of the DNA double helix structure.

The back-mapping procedure (*cgtoaa* module) makes use of HADDOCK’s versability in using distance restraints. During the initial CG mapping stage, atoms-to-bead restraints are defined, consisting of one distance restraint per bead 0Å (lower and upper distance defined as 0) between the geometric center of the atoms and the bead to which they belong. These restraints are used to morph the starting respective AA single structures onto the CG modelled complex. This back-mapping protocol consists of three main steps: (i) An initial fitting of the single AA structures onto their CG counterparts using the atoms-to-bead distance restraints, downscaling the intermolecular interactions to avoid clashes between molecules; (ii) short cycles of restrained minimization and molecular dynamics introducing conformational changes into the AA structures to match the conformation of the CG model, gradually increasing the interaction strength between molecules and; (iii) final clash removal through additional rounds of minimization and brief MD. During all those steps, the CG model is kept rigid and fixed in space, and both AA and CG models co-exist, but don’t see each other except through the effect of the CG to AA distance restraints. At the end of the protocol, the CG model is deleted. For more information, refer to the Methods section of ^23^.

### Docking protocols

All docking calculations were performed locally using HADDOCK3^11^. For comparison purposes, two docking workflows were performed, one with the standard all-atom representation using the united-atom OPLS force field^30^ and one in which docking and semi-flexible refinement are performed at the coarse-grained level with MARTINI2.2^21^, preceded by an AA to CG conversion step (*topocg*) and followed by the back-mapping step (*cgtoaa*).

The workflow is illustrated in Figure 1. A HADDOCK3 run always starts with the *topoaa* module, which defines the topology of the system and builds any missing atoms. In the CG pipeline, this is followed by the *topocg* module. Rigid body docking (*rigidbody*) is then performed. The topX models (with X=200 (default) or 400 in case of increased sampling) are then selected (*seletop*), followed by the *flexref* module, which performs a semi-flexible interface refinement of the models following a simulated annealing MD protocol in torsion angle space. In the CG pipeline, we then proceed to use the *cgtoaa* module to back-map the CG structures into their AA counterparts. The two final steps of the docking are the energy minimization (*emref*) module and the fraction of common contact-based clustering (*clustfcc*) module (with contacts identified with a cutoff distance of 7Å for the CG pipeline, instead of 5Å for AA). To analyze the quality and convergence of the models we added *caprieval* modules at various stages during the workflow (see Metrics for Evaluation of Model Quality below).

**Figure 1.**
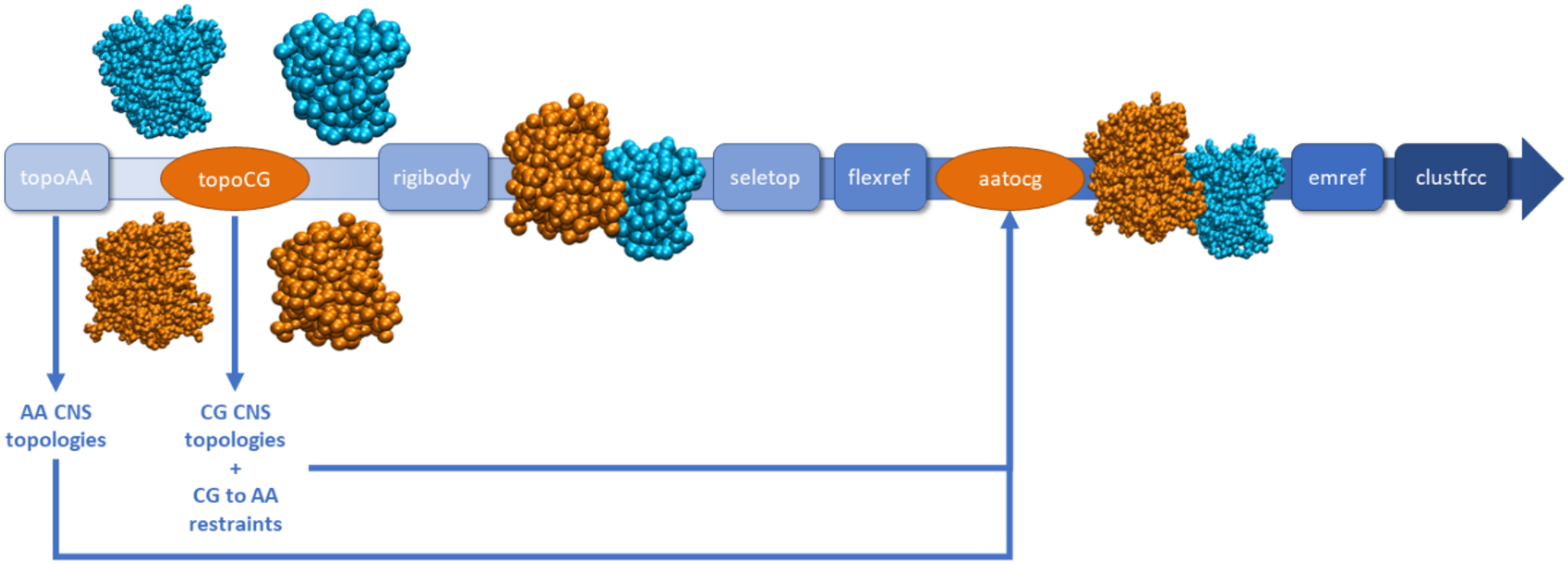
Coarse-grained HADDOCK3 docking workflow. AA stands for All-Atom, CG for Coarse-Grained. The two orange steps are only present in the CG workflow. *topoaa* defines the AA topology of the system and builds any missing atoms. *topocg* then maps the AA structures into their CG counterparts and defines the CG topology of the system. The *rigidbody* module performs the rigid body docking. *seletop* selects the top X models (X=200 in our benchmarks). The *flexref* module performs the semi-flexible interface refinement step, the *cgtoaa* module performs the back-mapping from CG representation to AA, and *emref*, the final energy minimization. The *clustfcc* module then performs a fraction of common contact-based clustering.

If a CG model is detected, the HADDOCK3 sampling (*rigidbody*) and refinement (*flexref, emre*) modules automatically adapts the non-bonded parameters to use a constant dielectric constant of 10, with a 14Å non-bonded cutoff and a switching function applied between 12Å and 14Å. For AA representations, a short cutoff of 8.5Å switched at 6.5Å is used (default in all HADDOCK versions) together with a distance-dependent dielectric constant (1/r). For protein-DNA docking, the dielectric constant is manually set to 78 by adding to the relevant modules (*rigidbody, flexref* and *emref)* of the workflow `epsilon=78`and`epsilon_cg=78`.

The HADDOCK3 workflows used for protein-protein and protein-DNA benchmarking are shown in Supplementary Material Figures SI-1 and SI-2 and available from the Git repositories of the benchmarking datasets.

### Protein-Protein and Protein-DNA Docking Benchmarks

To systematically test the performance of our CG protocol on protein-protein complexes, the Protein–Protein Docking Benchmark version 5.0^31^, containing 229 unbound-unbound cases, was used. A HADDOCK-ready version of this benchmark can be found on GitHub at https://github.com/haddocking/BM5-clean. Default sampling parameters were used (1000/200/200 for *rigidbody, flexref* and *emref*, respectively). The ambiguous interaction restraints (AIRs) guiding the docking were defined using the interfaces defined from the bound complex (true interface) by selecting all residues making intermolecular heavy atom contacts within a 5.0Å cutoff. These restraints model a rather ideal situation in which the interacting residues at the interface are known, even though their exact pairwise contacts are not, as would be the case when using experimental data such as NMR chemical shift perturbations or hydrogen/deuterium exchange mass spectrometry^32,33^.

For protein-DNA docking, we used 44 unbound-unbound cases from the protein-DNA benchmark^24,34^. A HADDOCK-ready version of the benchmark can be found on GitHub at https://github.com/haddocking/Prot-DNABenchmark. AIRs guiding the docking were taken from Van Dijk et al^34^ (available from the Git repo).

Note that, as the main purpose of this work was to re-implement and compare all-atom vs coarse-grained docking performance, we did not perform any ab-initio docking benchmarking.

### Scoring

HADDOCK’s scoring function incorporates the intermolecular electrostatic (E_elec_) and the van der Waals (E_vdW_) energies, an empirical desolvation energy term (E_desolv_)^35^, the buried surface area (E_BSA_), and the ambiguous interaction restraint energy (E_air_). The specific weights and contributions of these terms vary depending on the step of the docking protocol:

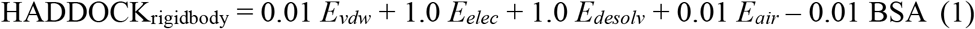

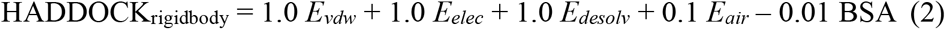

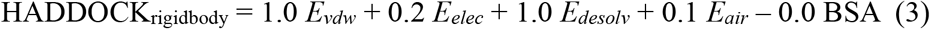

### Metrics for Evaluation of Model Quality

The quality of the generated models was evaluated using the *caprieval* module of HADDOCK3, which calculates the CAPRI metrics^36^, fraction of native contacts (FNAT), interface (i-RMSD) and ligand (l-RMSD) root-mean-square deviations from the reference experimental structure. FNAT is calculated using all heavy atom–heavy atom intermolecular contacts using a 5Å distance cutoff (CAPRI definition)^36^ for all-atom structures and a 7Å cutoff for coarse-grained representations, to account for the different resolution (based on contact analysis between the two representations, results not shown). Coarse-grained reference structures are provided to *caprieval* during the CG stage of the workflow. The i-RMSD is calculated on the interface backbone after superimposition on the interface residues, defined as those with any heavy atom within a 10Å distance of the partner protein. The l-RMSD is calculated on the ligand (usually the smallest molecule) after superimposition on the backbone atoms of the receptor (largest molecule). For both i-RMSD and l-RMSD, only backbone heavy atoms are considered (C_α_, C, N, O). Note that *caprieval* also returns the DockQ score^37^ (a combination of i-RMSD, l-RMSD and FNAT) and the interface ligand RMSD score (il-RMSD), which is useful for small molecules quality assessment. It is calculated by fitting the interface backbone of the receptor and calculating the RMSD of the ligand.

Based on these metrics, the quality of the docking poses is classified as:

- *High*: FNAT ≥ 0.5 and i-RMSD ≤ 1Å or l-RMSD ≤ 1Å (0.8 < DOCKQ),
- *Medium*: FNAT ≥ 0.3 and 1Å< i-RMSD ≤ 2 or 1Å < l-RMSD ≤ 5Å (0.49<DOCKQ<0.8),
- *Acceptable*: FNAT ≥ 0.1 and 2Å<i-RMSD≤4 or 5Å < l-RMSD ≤ 10Å (0.23 < DOCKQ < 0.49),
- *Near-Acceptable*: FNAT ≥ 0.1 and 4Å < i-RMSD ≤ 6Å,
- *Low quality*: FNAT < 0.1 or i-RMSD > 6Å or l-RMSD > 10Å.

### CG Integrative Modelling Examples with HADDOCK3

#### PFK filament

As a first application example, the phosphofructokinase-1 (PFK) filament in the R-state conformation was modeled (reference cryo-EM structure PDB ID: 8W2I^38^). This complex is composed of two R-state tetramers for a total of 5928 residues. To construct the filament from the unbound R-state conformation, the homotetramer from the cryo-EM structure of the R state (PDB ID: 8W2G^38^) was used as a starting point. The AIRs were defined from the interfaces extracted from the reference structure using the *ti* option from *haddock-restraints* (https://github.com/haddocking/haddock-restraints). Solvent accessible residues within a 3.9Å cutoff from the partner molecule were selected to define the ambiguous interaction restraints. The list of residues used for the docking is provided in Supplementary Table SI-1. Default sampling was used: 1000/200/200 for *rigidbody, flexref* and *emref*, respectively.

#### KaiC-KaiB complex

To model the KaiC:KaiB 1:6 complex, a single seven-molecule docking run was performed, targeting the CI domain of KaiC (see Snijder et al. ^39^ for details). The crystal structure of KaiC (PDB ID: 3DVL^40^), comprising 12 domains arranged in two six-membered rings, was used as the starting model. For KaiB, six copies of an NMR structure (PDB ID: 5JYT^41^) were used, which exhibits the same fold as in the reference complex, while the crystal structure^42^ originally used in Snijder et al. ^39^ corresponds to a fold-switched conformation. The cryo-EM structure of the KaiC:KaiB:KaiA complex was used as a reference (PDB ID:5N8Y^43^).

Residues previously identified by HDX-MS as solvent-protected in the CI domain of KaiC, and in KaiB^39^, were defined as active and passive residues, respectively, after filtering for solvent accessibility (relative residue solvent accessibility >50% as calculated with NACCESS^44^). A detailed list of these residues is provided in Supplementary Table SI-2.

KaiB monomers were restrained to an approximate C6 symmetry by defining three C2 symmetry pairs (B–E, C–F, D–G) and two C3 symmetry triplets (B–D–F and C–E–G).

In addition to the symmetry restraints, we established custom symmetry restraints which respect the C3 restraints, which ensures again the similarities between the 6 KaiB defined. As symmetry restraints were applied, sampling of 180° rotations during the rigid-body docking stage was disabled. The random removal of ambiguous restraints was removed as well. Furthermore, because of the number of subunits involved (7 in total), the sampling was increased to 10000/400/400 for *rigidbody, flexref*, and *emref*, respectively.

#### PRC1 Ubiquitation module bound to the Nucleosome

To model the Polycomb repressive complex 1 (PRC1) Ubiquitilation module bound to the Nucleosome, the unbound crystal structure of the enzymatic complex PRC1 was used (PDB code: 3RPG)^45^, and docked with the nucleosome particle (PDB code: 3LZ0). The default sampling values were used.

The docking was guided by interaction restraints reported in the literature available at the time of CAPRI Round 31 (Oct. 2014). These included a single unambiguous distance restraint (with an upper limit of 2Å) between the SG atom of the catalytic cysteine (Cys85) of PRC1 and the NZ atom of Lys118 on chain C of histone H2A, the ubiquitination site. In addition, mutagenesis data identifying key PRC1 residues (K62 of chain B, R64 of chain B, K97 of chain C, and R98 of chain C) as essential for nucleosome binding were incorporated^45,46^. AIRs were defined by assigning these PRC1 residues as active and all solvent-accessible histone residues on the side of the nucleosome where the catalytic cysteine is found as passive, filtered for solvent accessibility (the restraints and input structures were the same as in the protocol described in the HADDOCK2.4 server paper^47^). A complete list of the active and passive residues used to guide the docking is provided in Supplementary Table SI-3.

## RESULTS AND DISCUSSION

The docking protocol using MARTINI2, first implemented in HADDOCK2.4^23,24^, was ported to HADDOCK3^11^ through the addition of two modules: the *topocg* module, which maps the All-Atom (AA) structures to their Coarse-Grained (CG) counterparts, and the *cgtoaa* module, which performs the back-mapping of the CG structures into their AA representation. In the following section, we first discuss the performance of this implementation, comparing AA and CG docking results for the protein-protein and protein-DNA benchmarks. We then showcase the HADDOCK3 CG docking protocol by modelling three large complexes.

### Overall AA vs CG Performance for Protein-Protein and Protein-DNA docking

Two datasets were used for this comparison: the BM5 protein-protein benchmark^48^ (https://github.com/haddocking/BM5-clean), composed of 229 complexes, and the protein-DNA benchmark^49^ (https://github.com/haddocking/Prot-DNABenchmark), consisting of 44 complexes. The docking was performed from the unbound structures of each protein and driven by ambiguous interaction restraints derived from the real interface, which imitates an ideal scenario for HADDOCK3. The success rates at various stages throughout the docking were analyzed as the percentage of cases for which an acceptable or better model was predicted in the Top N models (Figure 2).

**Figure 2.**
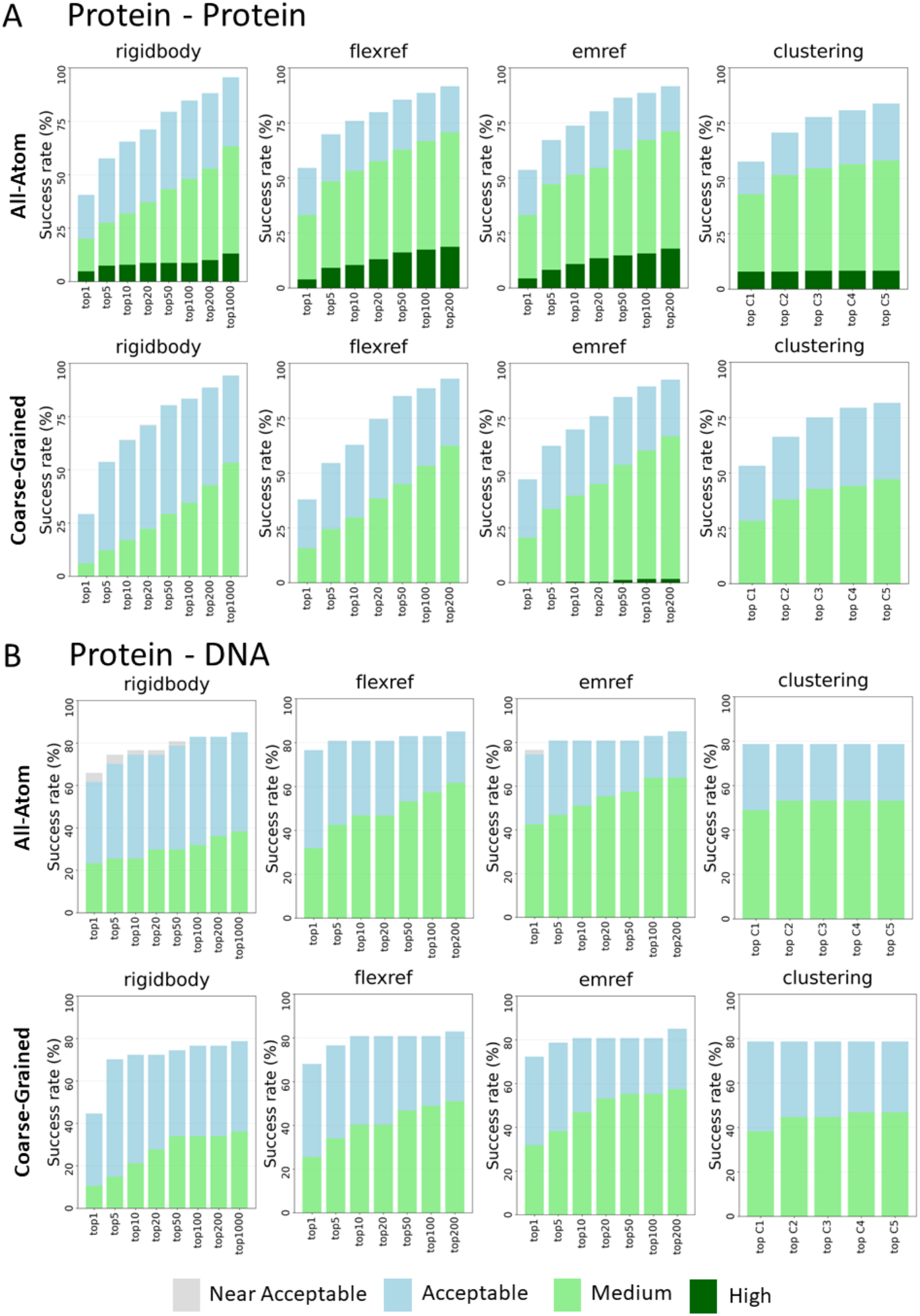
Performances of the AA and CG docking protocols in HADDOCK3. (**A**) Overall success rate (%) of the protein-protein benchmark as a function of the number of models considered, comparing AA and CG protocols. For the clustering analysis, the top four models from the highest-ranked clusters were considered, with “top C1” referring to the first-ranked cluster, “top C2” to the two highest-ranked clusters, and so on. (**B**) Same as (A) but for the protein-DNA benchmark.

The overall docking performance achieved at the all-atom level for the protein-protein benchmark is largely preserved in the coarse-grained approach for each step of the docking. After the final energy minimization step, the top 1 success rate for acceptable or higher quality is 53.7% and 47.2% for AA and CG protocols, respectively (Figure 2A). The success rate in the top 10 reaches 73.8% for AA and 69.9% for CG. In the top 5 clusters, the success rate reaches 83.8% and 81.6% for the AA and CG protocols, respectively. When evaluating model quality, the AA protocol generates higher quality models than the CG protocol (Figure 2A).

Protein-DNA benchmarking showed similar success rates between the CG and AA pipelines as well, as shown in Figure 1C. Interestingly, after the refinement stage (*emref*), the CG protocol achieved very similar overall performance as the AA approach, with top 1 success rates of 72.3% versus 74.5% and top 5 success rates of 78.7% versus 80.8%, respectively. This consistency is also observed after clustering, as seen for the top 1 ranked cluster, which reaches success rates of 78.7% for both protocols. However, similar to the protein-protein benchmark, the quality of the models is higher for the AA docking as measured by the success rates of medium quality models (Figure 2C). As the DNA was modelled as a perfect B-form helix, the amount of conformational changes between unbound and bound form prevents the obtention of high-quality models using a single docking round (a two-stage protocol leading to higher quality models was previously described^34^).

### Computational efficiency

Coarse-graining reduces the number of degrees of freedom in the system, thereby improving the computational efficiency of docking calculations. As shown in Figures 3A and 3C, coarse-grained docking runs (full workflows combining pre- and post-processing stages) are atwo-fold faster than their all-atom counterparts for both the protein–protein and protein–DNA benchmarks. A separate analysis of the *rigidbody* and *flexref* modules reveals an even larger performance gains, with speedups of 4.9- and 3.7-fold, respectively, for the protein–protein benchmark, and 5.0- and 3.8-fold, respectively, for the protein–DNA benchmark (Figures 3B and 3D). This difference in speed-up between specific stages and the overall workflow comes from the pre- and post-processing stages and, in particular, the *cgtoaa* back-mapping step in the CG protocol, which adds additional computation requirements to the full workflow.

**Figure 3.**
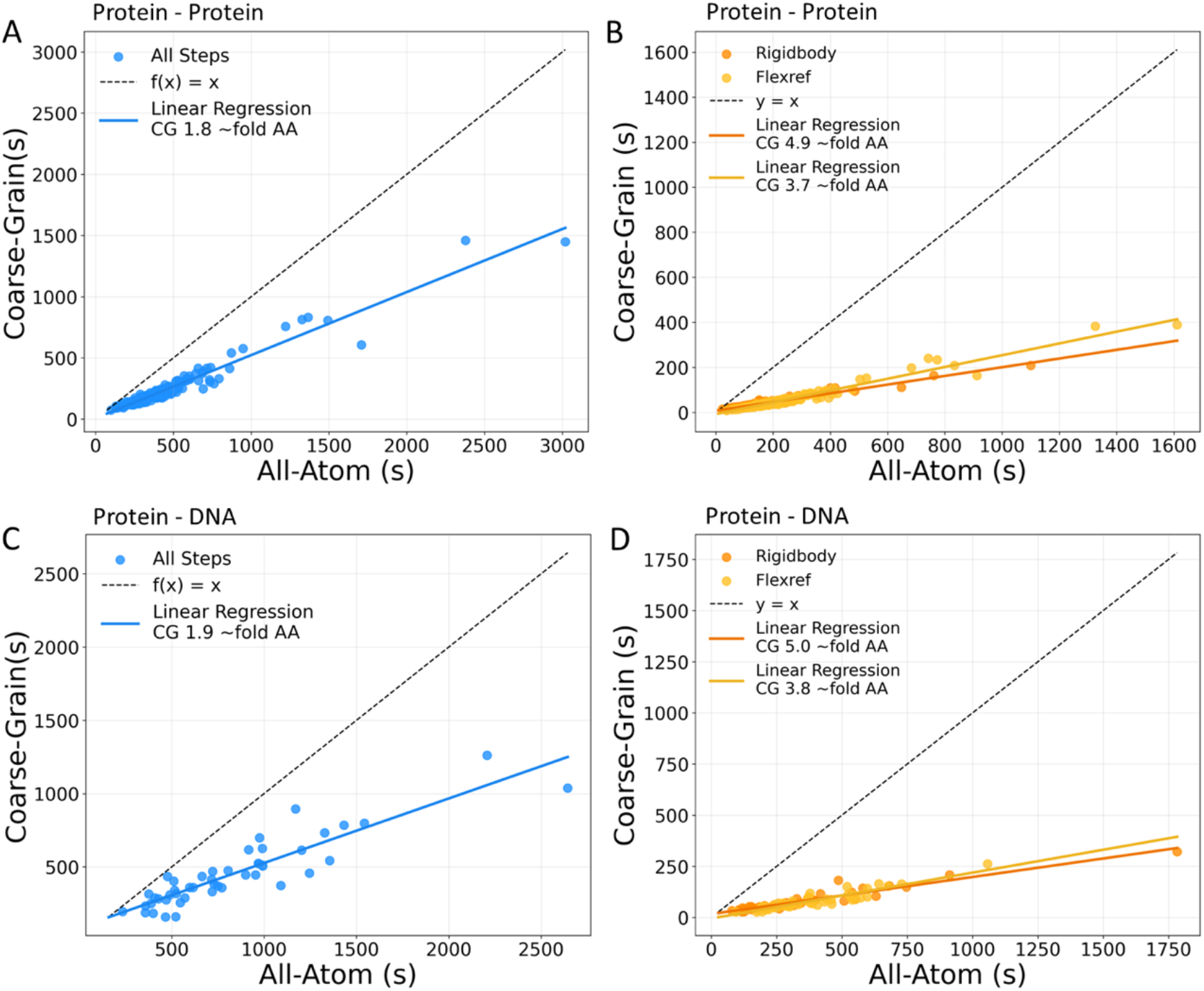
Correlation plot of total run time for CG versus AA simulations. (**A**) and (**B**): run times for the protein-protein benchmark, with all of the steps of the workflow (left), or *rigidbody* and *flexref* separated (right). (**C**) and (**D**): run times for the protein-DNA benchmark.

### CG Integrative Modelling Examples with HADDOCK3

#### The PFK filament: Modelling a dimer of large homotetrameric proteins

The capabilities of the HADDOCK3-CG approach were first illustrated using the phosphofructokinase-1 (PFK) test case. Two R-state PFK tetramers (PDB ID: 8W2G) were used to construct the filament structure. Interaction restraints were taken directly from the reference structure (PDB ID: 8W2I), mimicking an ideal docking case. The overall docking pipeline is similar to the protein-protein benchmark one, with sampling of 1000/200/200 for the rigid-body docking, flexible refinement, and energy minimization respectively.

The best-ranked model after the last energy minimization step was analyzed. The top-ranked prediction was classified as a medium-quality model according to CAPRI criteria, exhibiting an i-RMSD of 1.21Å, fraction of native contacts of 0.85 and DockQ of 0.71. The model aligned onto the reference structure is shown in Figure 4A. Notably, all models generated at the end of the docking run fall within the medium-quality category and cluster into a single cluster, highlighting the robustness and consistency of the HADDOCK3-CG protocol for this test case (using ideal restraints). All cluster statistics are reported in Supplementary Table 4.

**Figure 4.**
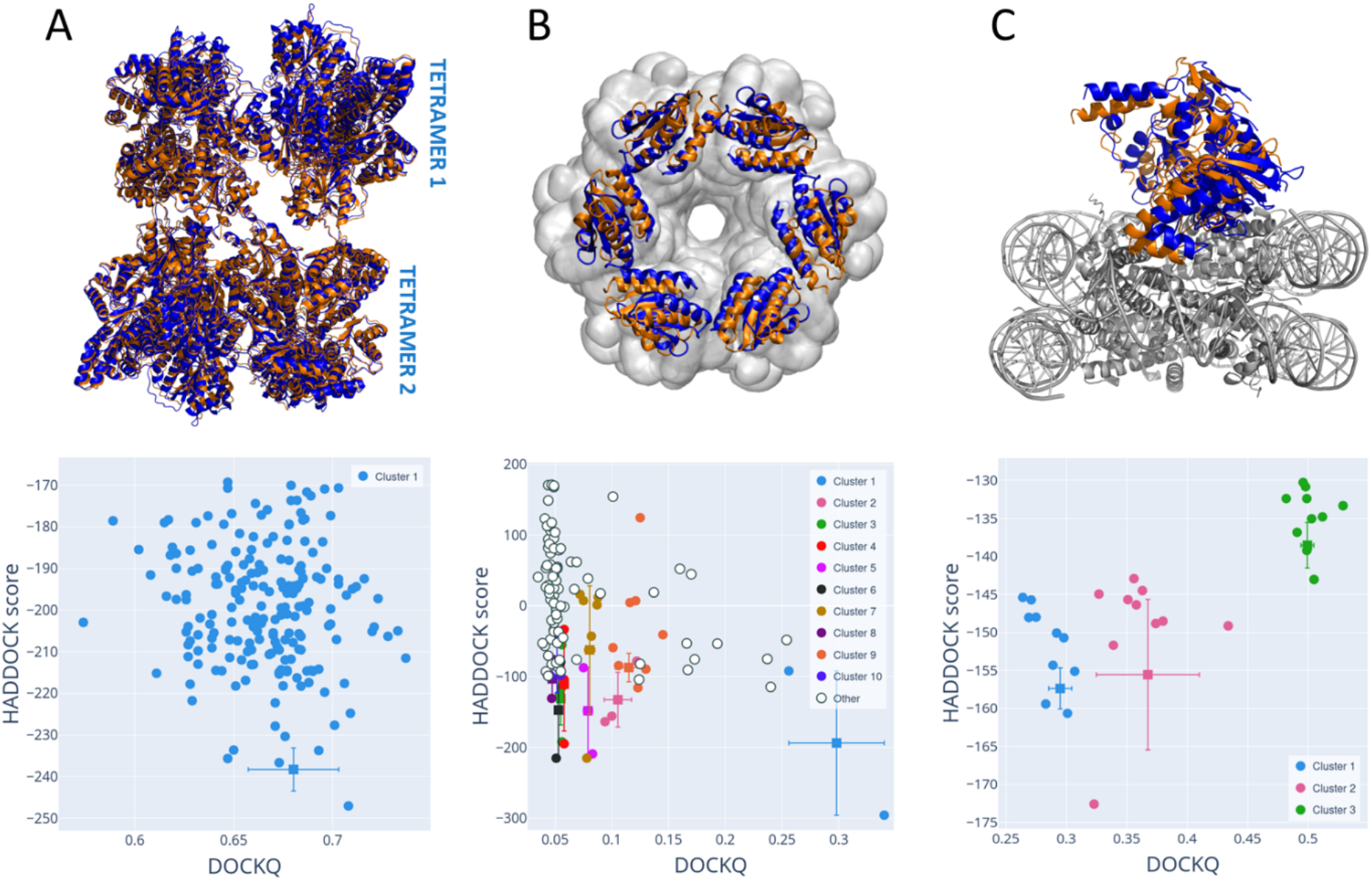
Structure comparison of top-ranking models predicted by HADDOCK3-CG. **(A)** Top 1 ranked model of PFK R-state filament (orange) superimposed with the reference (blue, PDB ID: 8W2L). On the second panel, the DockQ of each model after the clustering phase is shown as a function of the HADDOCK score. **(B)** Best CG model of KAiC-KaiB (orange) superimposed onto the cryo-EM model (blue, PDB ID: 5N8Y). On the second panel, the DockQ of each model after the clustering phase is shown as a function of the HADDOCK score. **(C)** First-ranked (orange) CG model of PRC1 superimposed onto the experimental crystal structure (blue, PDB ID: 4R8P). On the second panel, the DockQ of each model after the clustering phase is shown as a function of the HADDOCK score.

#### KaiC-KaiB: Symmetry-Restrained Docking of a Multi-Component Protein Assembly

The heptameric KaiC:KaiB (1:6) complex was modeled using simultaneous seven-body docking guided by the same hydrogen–deuterium exchange MS data used in a previous work^39^. KaiC consists of two hexamers of CI and CII domains, respectively^50,51^. It binds six KaiB monomers^52^. Earlier studies initially suggested two possible binding modes: CI or CII, and the initial modelling pointed to the CII binding mode^39^ based on an analysis of MS collision cross-section data. Subsequent cryo-EM data^43^, however, revealed that KaiB binds to the CI domain in a conformation corresponding to the NMR structure of KaiB (PDB ID: 5JYT) (which differs from the first available crystal structure of KaiB, showing a fold switch^41^). The CI vs CII binding analysis was addressed in the previous implementation of CG in HADDOCK2.4^23^, demonstrating that docking with the correct KaiB conformation was able to identify the correct binding mode. Here, as an illustration of the HADDOCK3-CG protocol, we focus exclusively on the CI binding mode, which was modeled using the full KaiC structure and six copies of the binding-competent KaiB conformation. The same docking procedure was applied as in our previous work^23^, with symmetry restraints approximating C6 symmetry between the six KaiB subunits by defining combined C3 and C2 relationships. In addition to the symmetry restraints, we established custom symmetry restraints which respect the C3 restraints, which ensures again the similarities between the 6 KaiB defined.

Analysis of the clustering results revealed that the near-native solutions were primarily found in the first-ranked cluster. In fact, the first-ranked structure (Figure 4B) was classified as an acceptable-quality model according to CAPRI criteria with an i-RMSD of 5.0Å, a FNAT of 0.37, l-RMSD of 7.30Å, and DockQ of 0.34. The corresponding metrics for all clusters are reported in Table SI-5.

#### Coarse-Grained modeling of the PRC1 Ubiquitination Module in Complex with the Nucleosome

The Polycomb repressive complex 1 (PRC1), which plays a central role in transcriptional repression during development by catalyzing histone ubiquitylation^46^, was used here as our third example to illustrate coarse-grained docking. Using interaction restraints derived from the literature (see Materials and Methods), the PRC1–nucleosome core particle complex was modeled with the MARTINI coarse-grained implementation in HADDOCK3 and the resulting models were evaluated against the available crystal structure of the complex (PDB ID: 4RP8^28^).

Based on the HADDOCK score, the top-ranked model (Figure 4C) from the first cluster is of acceptable quality with a DockQ of 0.32, FNAT of 0.29, l-RMSD of 7.8Å, and i-RMSD of 5.5Å. The best model of medium quality and with a DockQ of 0.529 was identified in the third-ranked cluster (6^th^ rank). The top three clusters are, however, all of acceptable quality or higher.

Cluster statistics are shown in Supplementary Table 6, showing the first 3 clusters to be in the acceptable to medium quality range. After energy minimization, 196 structures out of 200 are of acceptable or higher quality (27 medium quality). These results indicate that the coarse-grained docking protocol can successfully identify near-native PRC1–nucleosome configurations using literature-derived interaction restraints.

## CONCLUSIONS

In this work, we implemented the coarse-grained MARTINI force-field into HADDOCK3 and evaluated its performance, aiming at extending integrative docking with HADDOCK3 to large and complex biomolecular systems. Benchmarking on protein–protein and protein–DNA datasets demonstrates that the coarse-grained protocol largely preserves the docking success rates observed with all-atom calculations. Importantly, the reduced representation accelerates simulations by decreasing the number of interactions, enabling faster sampling and making the docking of large assemblies more computationally feasible. The applicability of the CG protocol in HADDOCK3 was further illustrated with three example cases—a phosphofructokinase-1 filament, the heptameric KaiC:KaiB (1:6) complex, and the PRC1– nucleosome core particle complex—where near-native models were successfully generated using experimentally derived interaction restraints. Together, these results highlight the usage of CG representation of molecules in HADDOCK3 as a robust and efficient framework for integrative modelling of large biomolecular assemblies.

## DATA AVAILABILITY

The protein-protein and protein-DNA benchmarking data (in HADDOCK-ready format) are available from https://github.com/haddocking/BM5-clean and https://github.com/haddocking/Prot-DNABenchmark respectively. All input data and results for the three large complexes used to illustrate the coarse-grained protocols are available from Zenodo (DOI: 10.5281/zenodo.19551728). The HADDOCK3 software can be obtained from https://github.com/haddocking/haddock3.

## ACKNOWLEGMENTS

The authors acknowledge funding from the European High Performance Computing Joint Undertaking projects BioExcel (Center of Excellence for Computational Biomolecular Research) (101093290).

