## Supplementary material for "Integrating the MARTINI2 Coarse-Grained Force Field into HADDOCK3 for Faster Modelling of Large Biomolecular Complexes"

### HADDOCK3 CG implementation

**Figure SI-1:** Detailed workflow for the protein-protein benchmark for both AA (A) and CG protocols (B).

A

```
run_dir = "run_AA_prot_prot"
molecules = [
    "data/receptor.pdb",
    "data/ligand.pdb",
]

[topoaa]

[rigidbody]
ambig_fname = "data/ambig_restraints.tbl"
ligand_top_fname = "data/ligand.top"
ligand_param_fname = "data/ligand.param"

[caprieval]
reference_fname = "data/target.pdb"

[seletop]

[flexref]
tolerance = 95
ambig_fname = "data/ambig_restraints.tbl"
unambig_fname = "data/unambig_restraints.tbl"
ligand_top_fname = "data/ligand.top"
ligand_param_fname = "data/ligand.param"

[caprieval]
reference_fname = "data/target.pdb"

[emref]
ambig_fname = "data/ambig_restraints.tbl"
unambig_fname = "data/unambig_restraints.tbl"
ligand_top_fname = "data/ligand.top"
ligand_param_fname = "data/ligand.param"

[caprieval]
reference_fname = "data/target.pdb"

[clustfcc]

[seletopclusts]

[caprieval]
reference_fname = "data/target.pdb"
```

B

```
run_dir = "run_CG_prot_prot"
molecules = [
    "data/receptor.pdb",
    "data/ligand.pdb",
]

[topoaa]

[topocg]
cgffversion = "martini2"

[rigidbody]
ambig_fname = "data/ambig_restraints.tbl"
ligand_top_fname = "data/ligand.top"
ligand_param_fname = "data/ligand.param"

[caprieval]
reference_fname = "data/target.pdb"

[seletop]

[flexref]
tolerance = 95
ambig_fname = "data/ambig_restraints.tbl"
unambig_fname = "data/unambig_restraints.tbl"
ligand_top_fname = "data/ligand.top"
ligand_param_fname = "data/ligand.param"

[caprieval]
reference_fname = "data/target.pdb"

[emref]
ambig_fname = "data/ambig_restraints.tbl"
unambig_fname = "data/unambig_restraints.tbl"
ligand_top_fname = "data/ligand.top"
ligand_param_fname = "data/ligand.param"

[cgtoaa]
ligand_top_fname = "data/ligand.top"
ligand_param_fname = "data/ligand.param"

[caprieval]
reference_fname = "data/target.pdb"

[clustfcc]

[seletopclusts]

[caprieval]
reference_fname = "data/target.pdb"
```

**Figure SI-2**: Detailed workflow for the protein-DNA benchmark for both AA (A) and CG protocols (B).

A

```
run_dir = "run_AA_prot_DNA"
molecules = [
    "data/receptor.pdb",
    "data/ligand.pdb",
]

[topoaa]

[rigidbody]
ambig_fname = "data/ambig_restraints.tbl"
ligand_top_fname = "data/ligand.top"
ligand_param_fname = "data/ligand.param"
epsilon = 78
dielec = "cdie"
w_desolv = 0

[caprieval]
reference_fname = "data/target.pdb"

[seletop]

[flexref]
tolerance = 95
ambig_fname = "data/ambig_restraints.tbl"
unambig_fname = "data/unambig_restraints.tbl"
ligand_top_fname = "data/ligand.top"
ligand_param_fname = "data/ligand.param"
epsilon = 78
dielec = "cdie"
dnarest_on = true
w_desolv = 0

[caprieval]
reference_fname = "data/target.pdb"

[emref]
ambig_fname = "data/ambig_restraints.tbl"
unambig_fname = "data/unambig_restraints.tbl"
ligand_top_fname = "data/ligand.top"
ligand_param_fname = "data/ligand.param"
dnarest_on = true
w_desolv = 0

[caprieval]
reference_fname = "data/target.pdb"

[clustfcc]

[seletopclusts]

[caprieval]
reference_fname = "data/target.pdb"
```

B

```
run_dir = "run_CG_prot_DNA"
molecules = [
    "data/receptor.pdb",
    "data/ligand.pdb",
]

[topoaa]

[topocg]
cgffversion = "martini2"

[rigidbody]
ambig_fname = "data/ambig_restraints.tbl"
ligand_top_fname = "data/ligand.top"
ligand_param_fname = "data/ligand.param"
epsilon = 78
dielec = "cdie"
w_desolv = 0

[caprieval]
reference_fname = "data/target.pdb"

[seletop]

[flexref]
tolerance = 95
ambig_fname = "data/ambig_restraints.tbl"
unambig_fname = "data/unambig_restraints.tbl"
ligand_top_fname = "data/ligand.top"
ligand_param_fname = "data/ligand.param"
epsilon = 78
dielec = "cdie"
dnarest_on = true
w_desolv = 0

[caprieval]
reference_fname = "data/target.pdb"

[emref]
ambig_fname = "data/ambig_restraints.tbl"
unambig_fname = "data/unambig_restraints.tbl"
ligand_top_fname = "data/ligand.top"
ligand_param_fname = "data/ligand.param"
dnarest_on = true
w_desolv = 0

[cgtoaa]
ligand_top_fname = "data/ligand.top"
ligand_param_fname = "data/ligand.param"

[caprieval]
reference_fname = "data/target.pdb"

[clustfcc]

[seletopclusts]

[caprieval]
reference_fname = "data/target.pdb"
```

**Table SI-1**: Detailed list of residues used as interaction restraints in the docking process for the PFK filament. Note that the numbering here is that of the experimental structure (PDB ID: 8W2I). For docking the numbering was adapted to that of the pre-processed input structures with the monomers merged into a single chain with non-overlapping numbering.

| Protein | Ambiguous Restraints Type | Residues |
|---|---|---|
| **R-state tetramer 1** | Active | chain B: Cys170, Gly171, Leu346, Pro347, Glu350, Met354, Leu373, Arg374, Arg485, Arg511, Gly512, Arg513, Tyr514, Glu515, Glu516, Cys518, Glu692, Arg695, Lys696, Arg698, Phe700, Asn702, Ala703<br>chain D: Tyr694, Arg695, Lys696, Gly697 |
| **R-state tetramer 2** | Active | chain A: Tyr694, Arg695, Lys696, Gly697<br>chain C: Cys170, Gly171, Leu346, Pro347, Glu350, Met354, Leu373, Arg374, Arg485, Arg511, Gly512, Arg513, Tyr514, Glu515, Glu516, Cys518, Glu692, Arg695, Lys696, Arg698, Phe700, Asn702, Ala703 |

**Table SI-2**: Detailed list of residues used as interaction restraints in the docking process for the KaiC-KaiB complex. Note that the numbering here is that of the experimental structures. As KaiC is an hexamer, the numbering of the six copies was shifted by 1000,2000,3000,4000,5000 and 6000 respectively and restraints were defined for each copy, targeting each a specific KaiB monomer. For KaiB, the residues identified from H/D data and their surface neighbors were both treated as passive residues in HADDOCK.

| Protein | Ambiguous Restraints Type | Residues |
|---|---|---|
| **KaiC (CI)** | Active | Gly101, Ile105, Asp107, Ala108, Pro110, Asp111, Pro112, Glu113, Gly114, Gln115, Glu116, Val117, Val118, Gly119, Asp122, Ser124, Ala125 |
| **KaiB** | Passive (from H/D data) | Thr7, Asn17, Thr18, Pro19, Glu33, Glu35, Gly38, Lys43, Leu48, Lys49, Pro51, Gln52, Glu55, Glu56, Lys58, Leu60, Pro70, Pro71, Pro72, Val73, Arg74, Ile77, Ser81, Asn82, Glu84, Lys85, Ile88 |
| **KaiB** | Passive (surface neighbors) | 9, 11, 15, 16, 20, 22, 23, 25, 29, 30, 34, 37, 39, 40, 41, 42, 44, 45, 46, 50, 53, 57, 59, 61, 62, 63, 64, 68, 69, 75, 76, 78, 79, 83, 87, 89, 91, 92, 94, 96, 97 |

**Table SI-3**: Detailed list of residues used as interaction restraints in the docking process. Residue numbering for the nucleosome follows a HADDOCK-adapted scheme, in which residue indices were offset by 200 for each chain (chains 1 to 10) to ensure unique numbering.

| Protein | Ambiguous Restraints Type | Residues |
|---|---|---|
| **PRC1 (PDB: 3RPG)** | Active | Lys62 (chain B), Arg64 (chain B), Lys97 (chain C), Arg98 (chain C) |
| **Nucleosome (PDB: 3LZ0)** | Passive | 39, 40, 41, 43, 44, 45, 49, 50, 53, 85, 88, 92, 94, 95, 96, 97, 250, 251, 252, 253, 254, 256, 257, 260, 261, 264, 281, 282, 285, 289, 292, 293, 296, 297, 300, 309, 500, 503, 506, 529, 530, 548, 552, 555, 556, 557, 558, 560, 572, 573, 575, 578, 579, 593, 594, 595, 597, 598, 600, 601, 602, 603, 762, 766, 767, 769, 770, 775, 779, 809, 810, 811, 814, 818, 823, 824, 827, 828, 831, 832, 834, 835, 838, 839, 841, 842, 843, 844, 1014, 1018, 1077 |

**Table SI-4:** Cluster-based statistics for the PFK filament. HADDOCK score single terms are averaged over the top 4 members of each cluster. Clusters are ordered according to the averaged HADDOCK score (a.u.). $E_{vdw}$: Lennard-Jones potential. $E_{elec}$: Coulomb potential. $E_{AIR}$: Ambiguous interaction restraints energy. $E_{desolv}$: Empirical desolvation score. BSA: Buried surface area. The CAPRI docking quality metrics are as follows: interface RMSD (i-RMSD), fraction of native contacts (FNAT), ligand RMSD (l-RMSD) and DOCKQ.

| **Cluster rank** | **1** |
|---|---|
| **Population** | 199 |
| **HADDOCK score** | -238.3 ± 5.2 |
| **$E_{vdw}$** | -54.5 ± 6.4 |
| **$E_{elec}$** | -1080.5 ± 71.8 |
| **$E_{AIR}$** | 37.1 ± 17.3 |
| **$E_{desolv}$** | 28.6 ± 6.8 |
| **BSA** | 2858 ± 120 |
| **i-RMSD (Å)** | 1.2 ± 0.02 |
| **l-RMSD (Å)** | 6.6 ± 1.0 |
| **FNAT** | 0.81 ± 0.03 |
| **DOCKQ** | 0.68 ± 0.02 |

**Table SI-5:** Cluster-based statistics for the KaiC-KaiB complex. HADDOCK score single terms are averaged over the top 4 members of each cluster. Clusters are ordered according to the averaged HADDOCK score (a.u.). $E_{vdw}$: Lennard-Jones potential. $E_{elec}$: Coulomb potential. $E_{AIR}$: Ambiguous interaction restraints energy. $E_{desolv}$: Empirical desolvation score. BSA: Buried surface area. The CAPRI docking quality metrics are as follows: interface RMSD (i-RMSD), fraction of native contacts (FNAT), ligand RMSD (l-RMSD) and DOCKQ.

| **Cluster rank** | **1** | **2** | **3** | **4** |
|---|---|---|---|---|
| **Population** | 2 | 3 | 6 | 3 |
| **HADDOCK score** | -193.7 ± 102.0 | -159.6 ± 3.9 | -158.6 ± 33.3 | -149.4 ± 45.3 |
| **$E_{vdw}$** | -216.4 ± 1.5 | -266.5 ± 14.9 | -311.8 ± 39.3 | -214.1 ± 14.4 |
| **$E_{elec}$** | -2374.9 ± 647.8 | -1979.3 ± 149.0 | -1569.0 ± 145.9 | -2292.7 ± 197.5 |
| **$E_{AIR}$** | 4800.6 ± 38.4 | 4790.8 ± 25.7 | 4569.3 ± 43.5 | 4948.6 ± 91.5 |
| **$E_{desolv}$** | 17.6 ± 30.0 | 23.7 ± 13.6 | 10.0 ± 18.8 | 28.4 ± 11.1 |
| **BSA** | 11002 ± 1166 | 12469 ± 91 | 12686 ± 859 | 10213 ± 447 |
| **i-RMSD (Å)** | 5.4 ± 0.4 | 10.4 | 17.9 ± 0.1 | 15.6 ± 0.1 |
| **l-RMSD (Å)** | 8.2 ± 0.9 | 14.8 ± 0.2 | 24.7 ± 0.4 | 21.1 ± 0.2 |
| **FNAT** | 0.3 ± 0.06 | 0.02 | 0.05 | 0.02 |
| **DOCKQ** | 0.3 ± 0.04 | 0.1 | 0.06 | 0.06 |

**Table SI-6:** Cluster-based statistics for the CG wokflow of the PRC1 complex. HADDOCK score and its components are reported as averages over the top 4 members of each cluster. The clusters are ordered according to the averaged HADDOCK score (a.u.). $E_{vdw}$: Lennard-Jones potential. $E_{elec}$: Coulomb potential. $E_{AIR}$: Ambiguous interaction restraints energy. $E_{desolv}$: Empirical desolvation score. BSA: Buried surface area. The CAPRI docking quality metrics are as follows: interface RMSD (i-RMSD), fraction of native contacts (FNAT), ligand RMSD (l-RMSD) and DOCKQ.

| **Cluster rank** | **1** | **2** | **3** |
|---|---|---|---|
| **Population** | 10 | 10 | 10 |
| **HADDOCK score** | -157.4 ± 2.7 | -155.6 ± 9.9 | -138.6 ± 3.0 |
| **$E_{vdw}$** | -25.5 ± 1.6 | -22.0 ± 5.7 | -8.8 ± 3.4 |
| **$E_{elec}$** | -897.2 ± 22.4 | -915.6 ± 31.2 | -894.6 ± 23.0 |
| **$E_{AIR}$** | 74.5 ± 13.9 | 79.8 ± 4.2 | 77.6 ± 11.8 |
| **$E_{desolv}$** | 40.1 ± 2.5 | 41.6 ± 2.3 | 41.5 ± 2.0 |
| **BSA** | 2168 ± 64 | 2210 ± 151 | 1906 ± 79 |
| **i-RMSD (Å)** | 5.9 ± 0.04 | 5.2 ± 0.2 | 4.8 ± 0.06 |
| **l-RMSD (Å)** | 7.2 ± 0.1 | 5.9 ± 0.9 | 3.75 ± 0.1 |
| **FNAT** | 0.24 ± 0.03 | 0.35 ± 0.06 | 0.57 ± 0.02 |
| **DOCKQ** | 0.30 ± 0.01 | 0.37 ± 0.04 | 0.50 ± 0.01 |
